# Allometric equation for the commonest palm in the Central Congo Peatlands, *Raphia laurentii* De Wild

**DOI:** 10.1101/2022.08.22.503797

**Authors:** Yannick Enock Bocko, Grace Jopaul Loubota Panzou, Greta Christina Dargie, Yeto Emmanuel Wenina Mampouya, Mackline Mbemba, Jean Joël Loumeto, Simon L. Lewis

## Abstract

The world’s largest tropical peatland lies in the central Congo Basin. *Raphia laurentii* De Wild. is a palm which forms dominant to mono-dominant stands across ca. 45 % of the peatland area. However, a lack of allometric equation for this canopy-forming trunkless palm with fronds up to 20 m long, means that this it is currently excluded from aboveground biomass (*AGB*) estimates for these peatland ecosystems. Here we develop an allometric equation for *R. laurentii*, by destructively sampling 90 *R. laurentii* stipes (across six mean petiole diameter size classes of: 2-4 cm, 4-5 cm, 5-6 cm, 6-7 cm, >8 cm) in a peat swamp forest, in the Likouala Department, Republic of the Congo. Prior to destructive sampling, the five parameters were measured for each stipe: diameter at stem base, mean diameter of the palm fronds, total diameter of the palm fronds (*TD*_*pf*_), total height and number of palm fronds. After destructive sampling and before weighing each individual was separated into the stem and the following palm frond categories: sheath, petiole, rachis and leaflets. We fitted a linear model relating *AGB* to each independent predictor variable separately to assess the best variable, finding that palm fronds represented at least 77 % of the total *AGB* in each diameter classes. We found that *TD*_*pf*_ was the best single predictor variable for *R. laurentii AGB*. We provide a series of equations, based on sampling *R. laurentii* with fronds ≥2 cm diameter or ≥5 cm diameters. The best allometric equation was: *AGB* = *Exp* (−2.691 + 1.425 × ln (*TD*_*pf*_) + 0.695 × *ln*(*H*) + 0.395 × ln (*WD*).. A monodominant 1 ha plot near the harvesting site had a palm AGB of 60.8 Mg ha^-1^, similar to the *AGB* of the trees in the same plot, at 86.2 Mg ha^-1^. Accounting for the *AGB* of palms is important, and can now be estimated using the allometric equations developed here.

## Introduction

Improved estimates of the carbon stocks and flows in tropical ecosystems is critical to estimating carbon loss from land-use change, and carbon uptake in remaining ecosystems [1]. For example, the implementation of REDD+ (reducing emissions from deforestation and forest degradation) in tropical countries is crucial for climate change mitigation [2]. Each country engaged in the REDD+ process must provide information on their REDD+ strategies and carbon reference levels [3]. However, information on reference levels requires accurate estimates of the biomass, and therefore the carbon stock, of the different forest types and carbon pools: vegetation (dicotyledon and monocotyledon), coarse woody debris, litter and soil.

For the countries of the Congo Basin establishing carbon reference levels is hindered by a scarcity of data. This is particularly true for swamp forest ecosystems. The peatlands of the Cuvette Centrale Congo Basin are estimated to cover 145,500 km^2^ and store 30.6 Pg C belowground in the peat [4]. However, *in situ* data on aboveground biomass from these peatland ecosystems is rare [4–6]. Furthermore, these utilise the allometric equations available to estimate the biomass of trees, but these were developed using data from terra firme forests. Additionally, across ca. 45% of these peatlands the palm species *Raphia laurentii* De Wild [4] forms dominant or monodominant stands, but at present there is no obvious allometric equation which can be applied to *R. laurentii*. Hence in past estimates of central Congo peat swamp aboveground biomass palms have been excluded, leading to a systematic bias and underestimation [4–6]. This neglect follows a pattern, as globally, the contribution of palms to forest biomass and productivity is a neglected area of study, and therefore a source of uncertainty in forest carbon stocks and dynamics [7]. This extends to African tropical forests, where *R. laurentii* is likely to be one of the most abundant palm species in the Congo Basin.

One potential solution to the lack of an allometric equation for *R. laurentii* is to use an allometric equation from a palm from elsewhere in the tropics, as the use of a dicot allometric model for the estimation of the above-ground biomass of a monocot species has been found to be highly inadequate [8,9]. Whilst some allometric equations have been developed for Amazonian [9,10] and Asian [11,12] palm species, these are morphologically dissimilar to the trunkless *R. laurentii*. Generally, in monocot species, using a species-specific allometric model is better than using a multispecies model [9]. Therefore, the development of an allometric equation for what is probably the dominant canopy plant in the world’s largest tropical peatland complex, *R. laurentii*, is a fundamental contribution to understanding the carbon cycle in these peatland ecosystems. Without this information to inform sustainable land management policies, the peatlands are at risk from inappropriate development, which could have serious consequences for the region [13].

By linking the aboveground biomass (AGB) of a species to certain physical parameters that can be measured in the field [14], allometric equations permit the AGB of a species to be estimated non-destructively. The physical parameters that can be used to establish an allometric equation are numerous and differ depending on the species. Diameter at breast height (DBH), wood density, height [15] and crown diameter [16] are, in descending order, the most important physical parameters in the estimation of the biomass of a dicotyledonous species. By contrast, for monocots, depending on the architectural type of the stem, number of palm fronds, dry mass fraction, wood density, DBH, diameter at the base of the stem, crown diameter, total height and stem height are often used as physical parameters [8,17,18]. For palms in particular, height and diameter (at the base or at breast height) are the most important physical parameters for estimating above-ground biomass, although it is unclear if this would apply to trunkless palms as well [11,18].

To address the lack of palm allometry for the commonest canopy plant in the central Congo peatlands, the aim of this study is to develop an allometric equation for the monocot species *R. laurentii*, in order to improve the biomass and aboveground carbon stock estimates of the peat swamp forests of the Congo Basin.

## Material and Methods

### Study area

Fieldwork for this study was carried out in February 2019 in the Likouala Department, Republic of the Congo. The study site is located near to the village of Bolembe (1.190288N and 17.84239E; Fig. 1) in a peat swamp forest which occupies an interfluvial basin, extending ∼60 km between the black-water Likouala-aux-herbes River and the white-water Ubangui River [19]. The mean annual temperature and precipitation are respectively 25.6 °C and 1556.5 mm [20]. The peatland is shallowly domed [19] and the vegetation is composed of two communities; one dominated by dicotyledons such as *Carapa procera* DC., *Symphonia globulifera* L. f., *Uapaca mole* Pax, *Grosera macranta* Pax, *Manilkara fouillonna* Aubrev. & Pellegr. and *Drypetes occidentalis* (Mull. Arg.) Hutch., termed hardwood-dominated peat swamp vegetation; the other by monocotyledons such as *Raphia laurentii, Aframomum angustifolium* (Sonn.) K. Schum. and *Pondanus candelabrum* P. Beauv., termed palm-dominated peat swamp vegetation [4,21,22], with a gradation between these two vegetation types often spanning many kilometres. The fieldwork was conducted in a palm-rich area of swamp forest overlying ∼2 m of peat (Fig 1).

**Fig 1.**
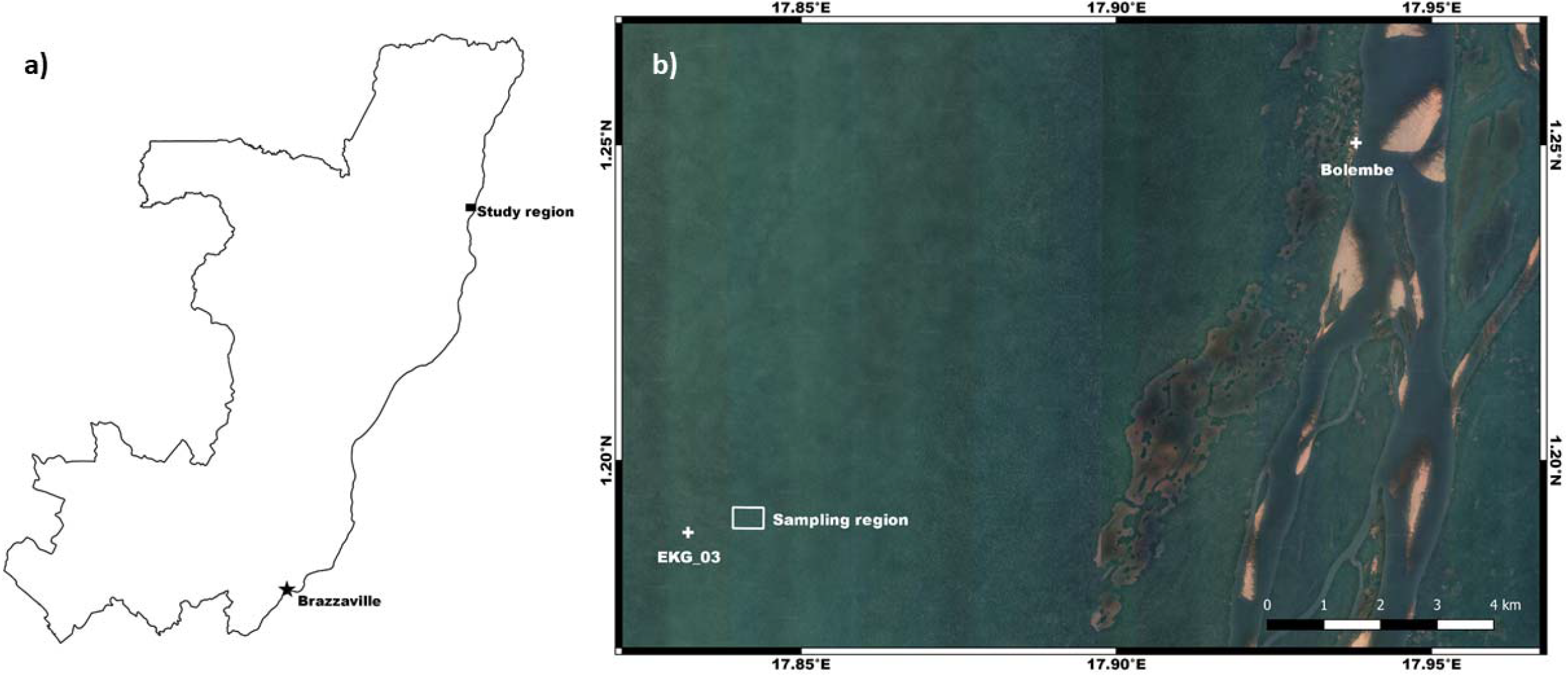
Location of the study area.

### Palm selection and sampling

Biomass allometry consists of predicting biomass from easily measurable variables [14]. Botanical characterisation of *R. laurentii* by [23] revealed it to be a cespitose palm (multiple shoots from the same root system) with 4 to 6 stipes (i.e. short trunks) up to 20 cm in diameter. A stipe can have a maximum of 10 palm fronds (each comprising a sheath of c. 1 m in length, a petiole c. 4 m in length and a rachis c. 3 to 4 m in length). Their architecture and their evolution in a compact clump makes it difficult to measure the diameter at the base of the stem or at 1.30 m from the ground. Thus, in order to select palms for destructive harvest across the size-range of the species to construct a robust allometric equation, palm frond diameter data were collected in five subplots (0.04 ha) of a permanently installed plot (1 ha), in a *R. laurentii* dominated forest, called EKG-03, located at (1.191998N, 17.84693E). in February 2019. The diameter of each palm frond was measured using callipers between the sheath and petiole at 1.30 m from the ground where possible or, when it was inaccessible at this point, slightly above 1.30 m. Petiole diameter varied between 2 and 11.5 cm, both between and within *R. laurentii* stipes. Based on the average petiole diameter for each palm stipe, we grouped the palms stipes into six mean petiole diameter classes: [2-4 cm], [4-5 cm], [5-6 cm], [6-7 cm], [7-8 cm] and ≥8 cm. For each mean petiole diameter class we then selected 15 stipes, each with at least three palm fronds, to be destructively samples outside the permanent plot, giving a total of 90 *R. laurentii* stipes.

Prior to felling, for each *R. laurentii* stipe the palm location was recorded with a global positioning system (GPS), the number of live palm fronds was counted [24] and the diameter at the base of the stem was taken approximately 13 cm from the ground [11]. The total height, defined as the length from the base of the stem to the apical meristem [10], was measured using a vertex. The diameter of each palm frond was measured on the petiole at 1.30 m from the ground or above 1.30 m at levels as low as possible on the petiole but avoiding the sheath, using a calliper.

### Destructive sampling and biomass data

Destructive sampling was carried out using a chainsaw (manufacturer: STIHL; model: MS780). We then measured the length of the felling section at the apical meristem as well as the height of the hinge, before separating each palm frond from the stem via the chainsaw at the base of the sheath. Each stipe of *R. laurentii* was then divided into stem (including the base of the sheath) and palm frond [18,25] components. Each palm frond was further subdivided into four parts: sheath, petiole, rachis and leaflets. The leaflets were detached from the rachis using a machete. The fresh mass (kg) of each palm frond part was weighed in the field using a hand-held balance and the lengths of the sheath, petiole and rachis were measured (Fig 2A). From each *R. laurentii* stipe, three palm fronds were selected and from these fronds five 15 cm samples [12] were collected from the middle of the sheath, the middle of the petiole, at the base of the rachis, the middle of the rachis and the top of the rachis (Fig 2B). A single sample was taken from the leaflets (a minimum of 400 g; Figure 2C), giving a total of 18 samples per *R. laurentii* stipe. For the stem, the length was measured, before we cut it into sections to allow the fresh mass (kg) to be weighed in the field using a hand-held balance with a 300 kg capacity. For each stipe, three discs of 5 cm were collected from the base, the middle and the top of each stem (Fig 2D), weighed with a 1 or 5 kg capacity balance before being sealed in plastic and transported to the laboratory.

**Fig 2.**
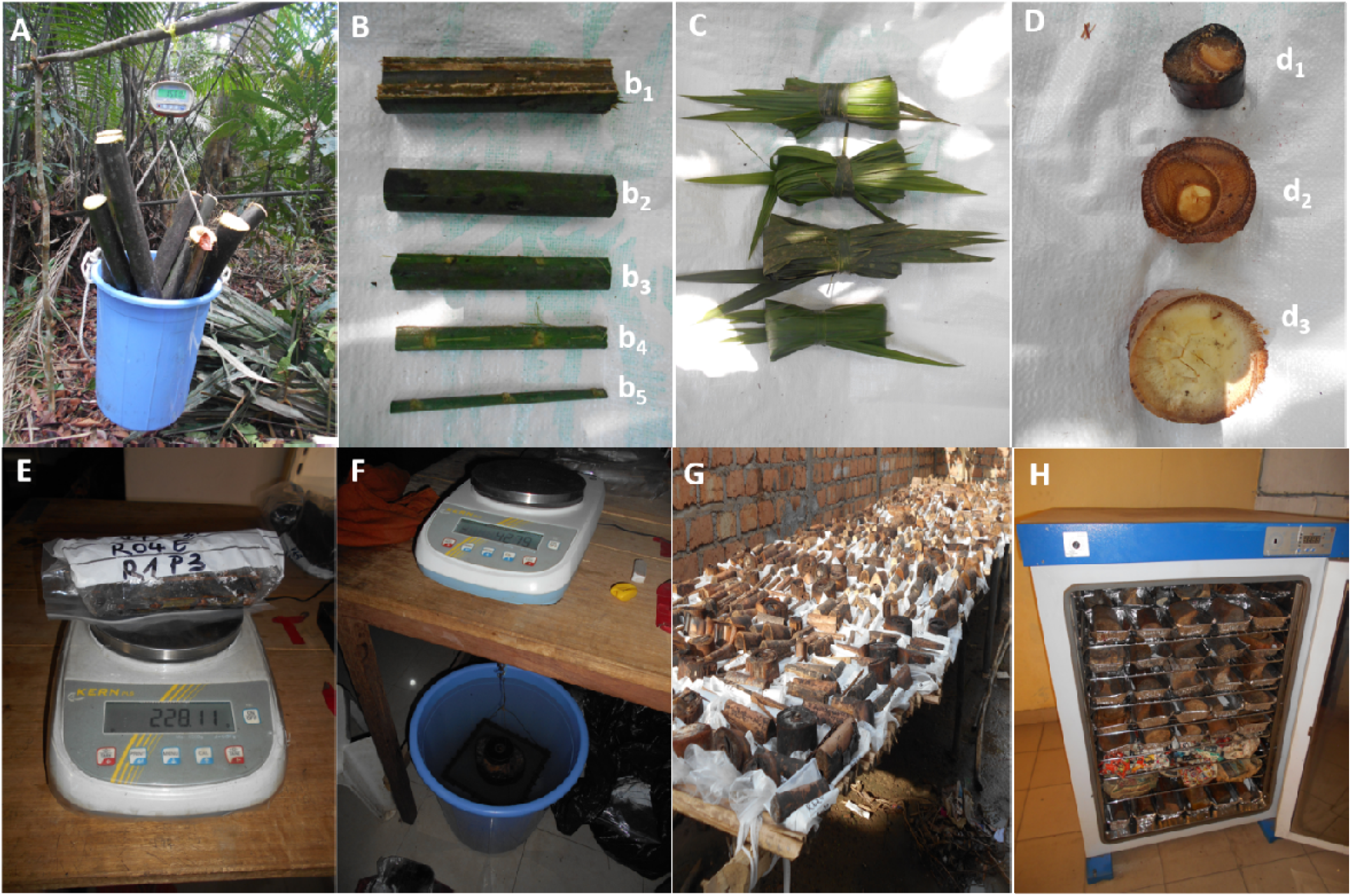
Illustration of field and laboratory data collection. Measurement of the fresh weight of part of the palm (A), samples taken from a palm (B), samples from leaflets (C), samples taken from the stem (D), measurement of fresh mass precision in the laboratory (E), measurement of green volume (F), sun drying (G) and oven drying (H). b_1_ : the middle of the sheath, b_2_ : the middle of the petiole, b_3_ : the base of the rachis, b_4_ : the middle of the rachis, b_5_ : the top of the rachis, d_1_ : the base of the stem, d_2_ : the middle of the stem and d_3_ : the top of the stem.

In the laboratory, all samples from the stem and palm fronds were weighed in their plastic bags using a suspended 6.2 kg capacity precision balance (Fig 2E). The mass (g) of each plastic bag was then subtracted. The volume (cm^3^) of each stem, sheath, petiole and rachis sample was measured by hydrostatic weighing (Fig 2F; [26]). All samples were air dried (Fig 2G) before being oven drying at 65 °C (for leaflets) and 101 °C (for the stem, sheath, petiole and rachis; Fig 2H). to constant mass (g) [26].

The wood density (WD) of each sample was calculated as dry mass divided by volume, in units of g cm^-3^ [27]. The mean dry mass/fresh mass ratio of each sample of the stem, sheaths, petioles, rachis and the leaflet was used to convert the fresh mass weighed in the field, to dry mass (kg). The total biomass of each *R. laurentii* stipe was calculated by summing the dry mass (kg) of the stem, sheaths, petioles, rachis and the leaflets.

### Allometric modelling and evaluating

The power function has been analysed and is often considered as the best functional form model for modelling aboveground biomass prediction in the Congo Basin tropical forest [28], and has been considered the best form for palms in the Amazon [10] and more broadly for tropical forest trees [29]. Therefore, the allometric model of the power function form suggested by Parresol [30] was selected for this study:

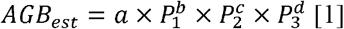

Where *AGB*_*est*_ is the aboveground biomass estimate of the palm stipe, *P*_*1*_, *P*_*2*_ and *P*_*3*_ are the independent predictor variables, *a* is the allometric coefficient and *b, c* and *d* are the allometric exponents. The variables were transformed into a natural logarithm, owing to the data being heteroscedastic [27,31], and the model was then written as follows:

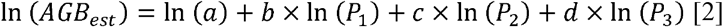

The systematic biases introduced by the logarithmic transformation were corrected using the correction factor defined below [15]:

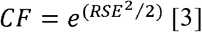

Where *CF* is the correction factor and 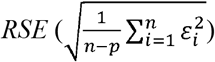 is the residual standard error of the linear model. Where *n* is the sample size, *p* is the total number of predictor variables in the model and □ is an error term. The independent predictor variables of the *AGB* were grouped into three types of parameters: coarseness parameter (diameter at stem base (*D*), mean diameter of the palm fronds (*MD*_*pf*_) and the total diameter of the palm fronds (*TD*_*pf*_)), a height parameter (total height (*H*) of the palm) and the parameters intrinsic to each palm stipe (wood density (*WD*) and number of palm fronds (*N*_*pf*_)). In order to select the best predictor variable, we fitted a linear model relating *AGB* to each independent predictor variable separately [32].

The tested linear model assumed *c = d =* 0 and Equation 2 was written as follows:

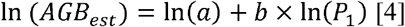

The best predictor variable was selected according the *RSE* and the Coefficient of Determination (*R*^*2*^). Afterwards, two types of a linear model were built. The first type assumed *d =* 0 when the best predictor variable (*P*_*1*_) was combined with one other predictor variable belonging to another type parameter (for example: *D with H, D with WD, MD*_*pf*_ with WD, *TD*_*pf*_ with *N*_*pf*_):

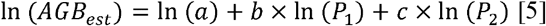

The second type of linear model included three predictor variables (the best with two other predictor variables) and assumed *b, c* and *d* were different to zero (equation 2).

## Data analysis

All statistical analyses were carried out with the R software package (http://www.r-project.org/). The *AIC*, the *RSE*, the *R*^*2*^_*ad*j_, the Roots Mean Square Error 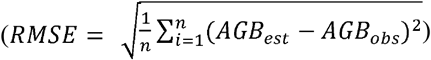 and the Bias prediction 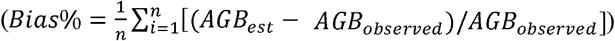were used for the evaluation of established models [26]. Linear models with higher *R*^*2*^_*ad*j_, smaller *AIC, RSE, Bias* and *RMSE*, and significant coefficients and *p-value* (p < 0.05) were selected as the best-fitted aboveground biomass allometric equations [33].

We also estimated the AGB of two 1 ha plots, the palm-dominated peat swamp forest plot EKG-03, and a hardwood dominated peat swamp forest, EKG-02, to show the impact of including R. *laurentii* in AGB estimates. All tress ≥10 cm diameter using standard protocols (Lewis et al. 2013). For palms the number of living fronds was counted and the diameter at the base of each stipe was measured where possible. To estimate the AGB of the trees, the equation of Chave et al. [29] using *D*, ρ and *E* was used to estimate the aboveground biomass of each tree, with ρ *from g*.*cm*^*-3*^ *and D from cm*. For the AGB of *R. laurentii* stipe we used one of the developed models, using palm frond number only. The quantification of the percentage difference in the biomass estimate of the two plots was done between the total biomass (trees and palm) and the tree biomass (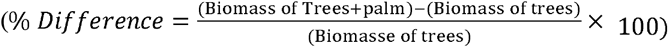)

## Results

### Aboveground biomass partitioning

The average number of fronds per stipe was 5.7±0.22 (mean±SE), with a range of 3– 11 fronds (n=90). The length of these fronds was 13.30±0.41 with a range of 5.13–21.20 m. The diameter at the base of the stipe was 21.67±0.73 with a range of 8.20 – 35.40 cm. The sum (total) of the petiole (frond) diameters was 37.57±2.67 cm, with a range of 7.20–125.40 cm and the average diameter of fronds per stipe was 6.01±0.22 cm with a range of 2.22–12.54 cm. The average of the total height per stipe was 13.30±0.41 m, with a range of 5.13–21.20 m. The minimum, maximum and average values of wood density, dry mass/fresh mass ratio and aboveground biomass of the individual parts of the 90 stipes of *R. laurentii* are presented in Table 1. The tissue density was low compared to trees, ranging from 0.15 to 0.38 g/cm^3^, with an average value of 0.24±0.003 g/cm^3^ and was approximately constant between the palm stem compartments and the palm fronds compartments (Table 1). The dry mass/fresh mass ratio varied from 0.16 to 0.55, with a mean value of 0.37±0.01 and was significantly different (P-value < 0.001) between the stem and the palm frond compartments.

**Table 1.**
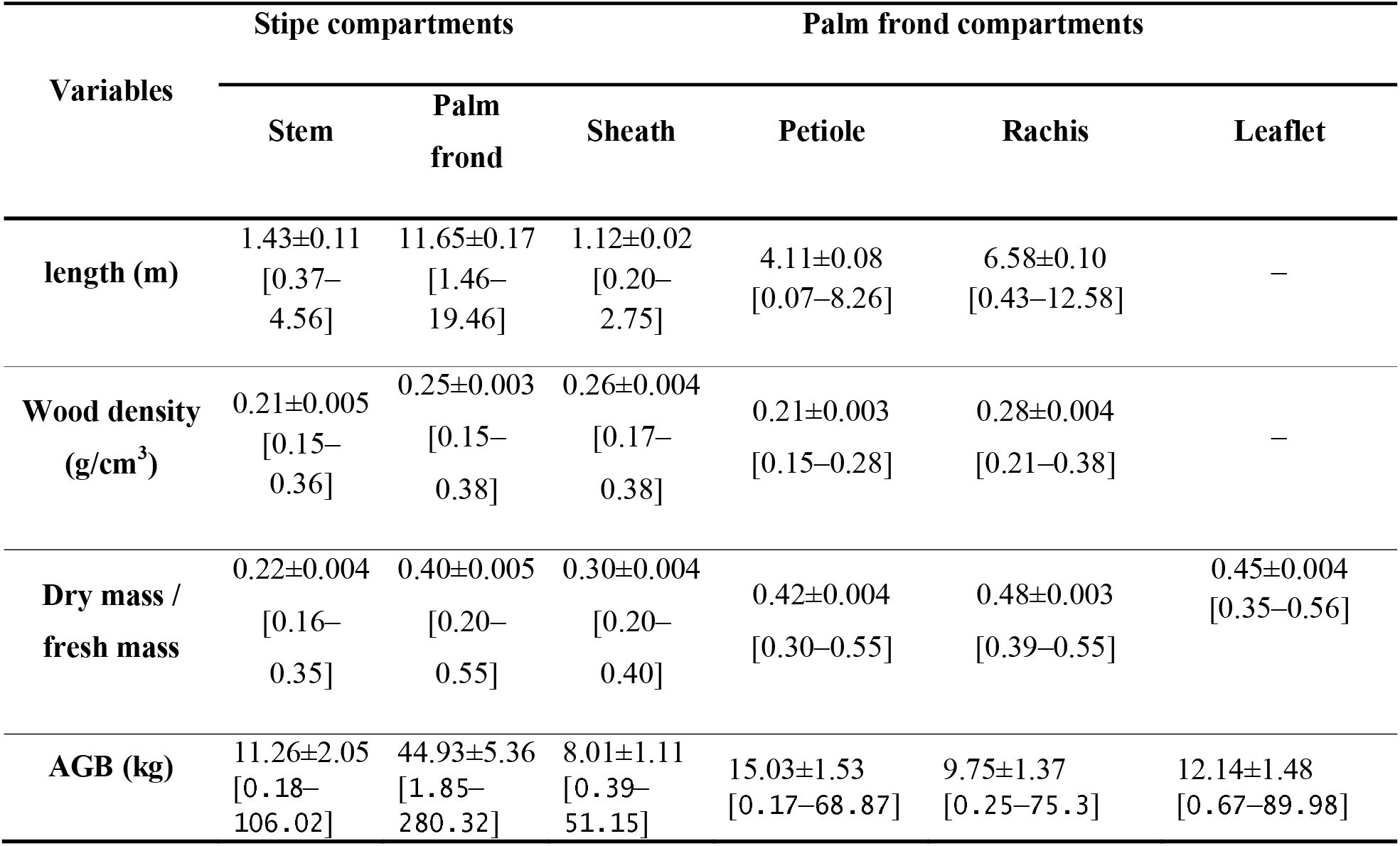
Physical characteristics (Mean±SE [Min–Max] value) of the 90 stipes of *R. laurentii* felled in the peat swamp forest, Likouala Department, Republic of the Congo.

The average aboveground biomass of the stem compartment (11.26±2.05 kg) was lower than that of the palm fronds (44.93±5.36). The allocation of biomass between each component (i.e., stem, sheath, petiole, rachis and leaflets) was approximately constant between mean petiole diameter classes (Fig 3). Palm frond compartments accounted for more than 77 % of the aboveground biomass in all mean petiole diameter classes (with the stem accounting for between 10–23 % of the total biomass). On average the petiole accounted for the largest proportion of biomass (30.17 %) followed by leaflets (25.17 %), rachis (16 %), stem (14.83 %) and sheath (13.83 %). While the proportion of rachis biomass increases with increasing mean stipe diameter classes, the proportion of leaflet biomass decreases (Fig 3).

**Fig 3.**
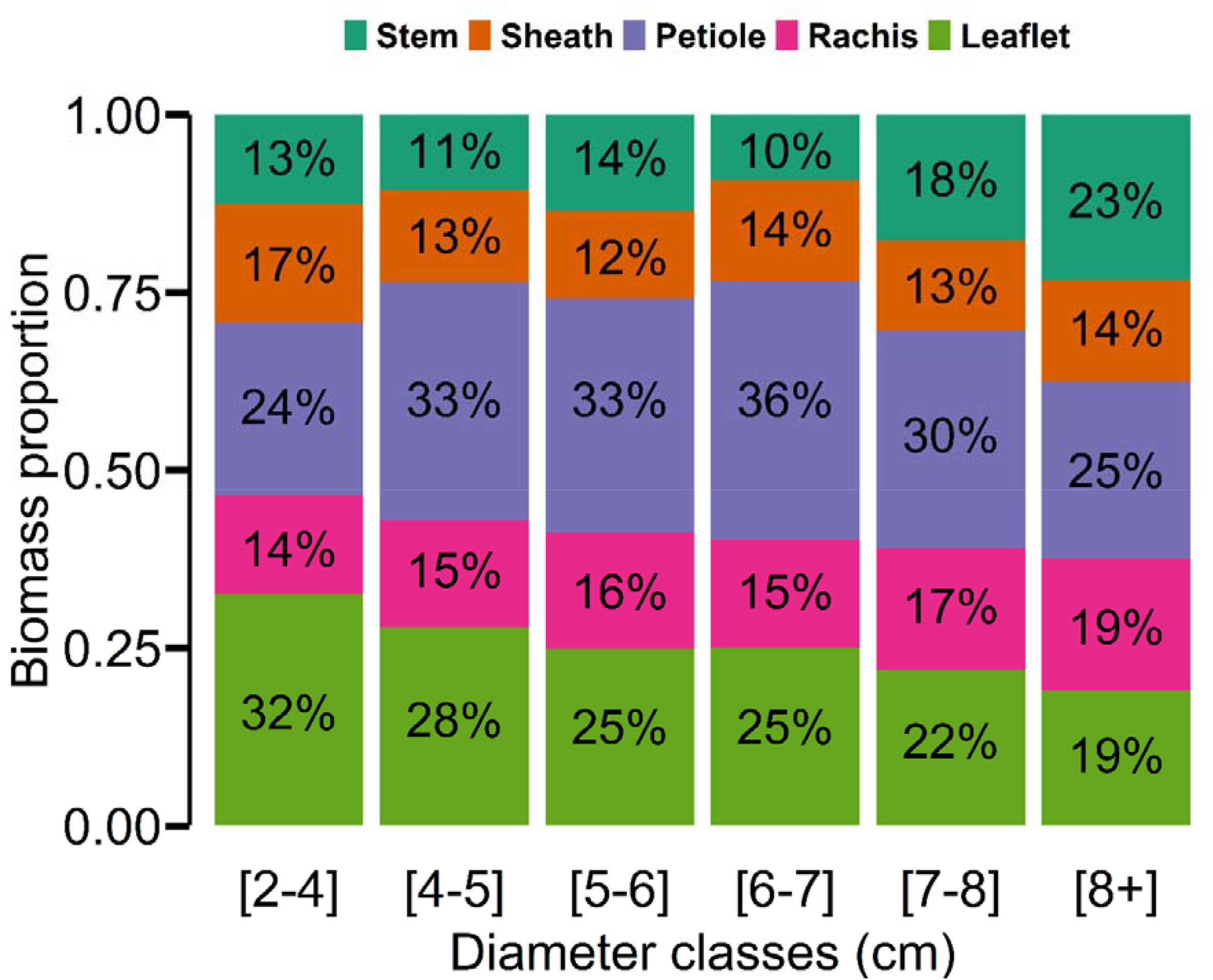
Aboveground biomass distribution among the stem, sheath, petiole, rachis and leaflet for each mean diameter class of the palm frond for the 90 individual *R. laurentii* sampled.

Palm frond *AGB* was significantly higher than the stem AGB in a stipe of *R. laurentii* (kruskal.test P-value < 0.001; Fig 4a). AGB increased with increasing mean petiole diameter class (Fig 4b).

**Fig 4.**
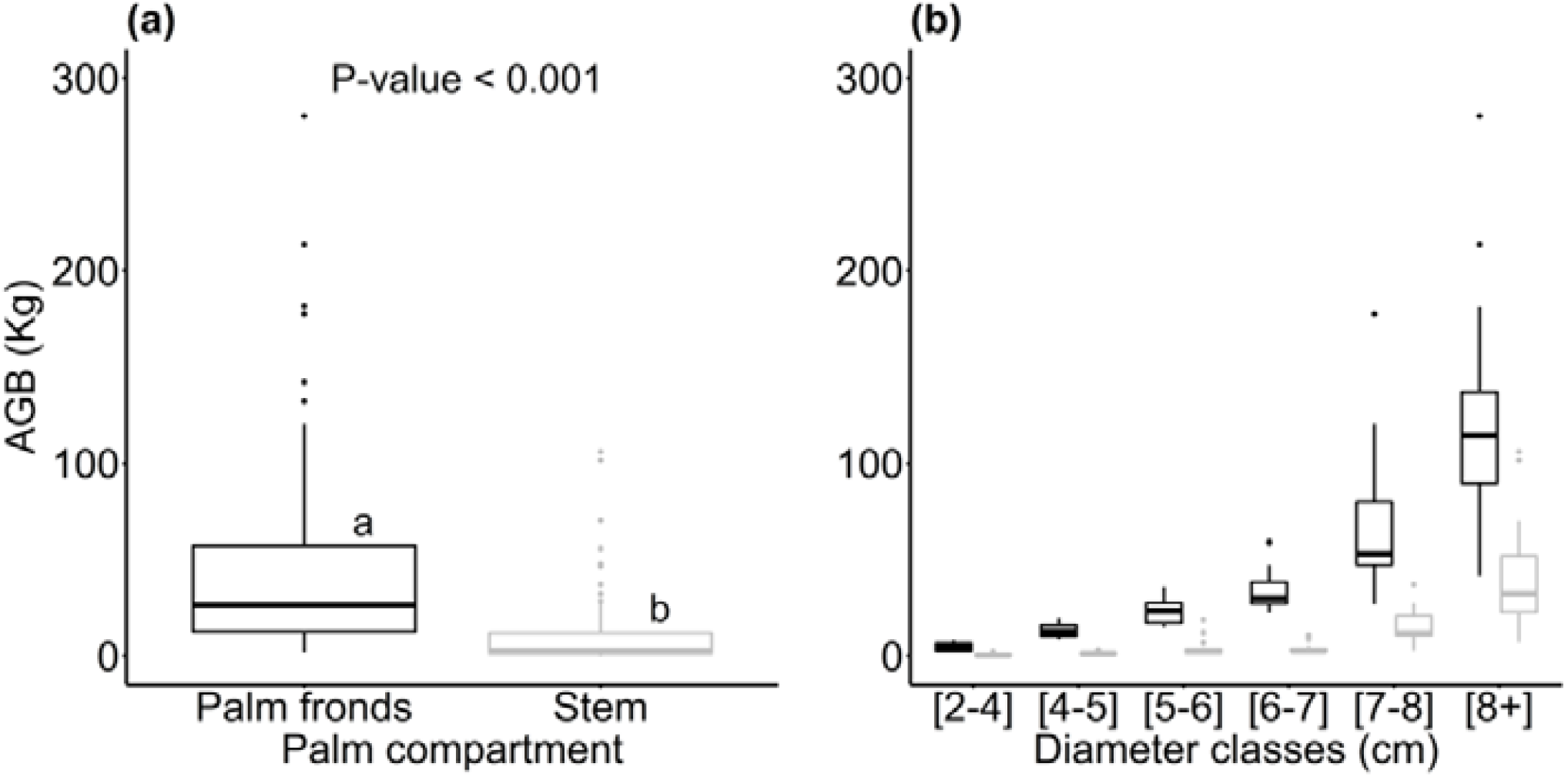
Variability of aboveground biomass between palm compartments (a) and diameter classes (b).

### Allometric equations for AGB estimation

In the analyses where only single predictor variables were used to estimate AGB and petioles ≥2 cm were included in the analysis, *TD*_*pf*_ was found to be the best single predictor, followed by *MD*_*pf*_, *D, H* and *N*_*pf*_ respectively, with all five variables showing a strong relationship with AGB (Fig 5a-e).

**Fig 5.**
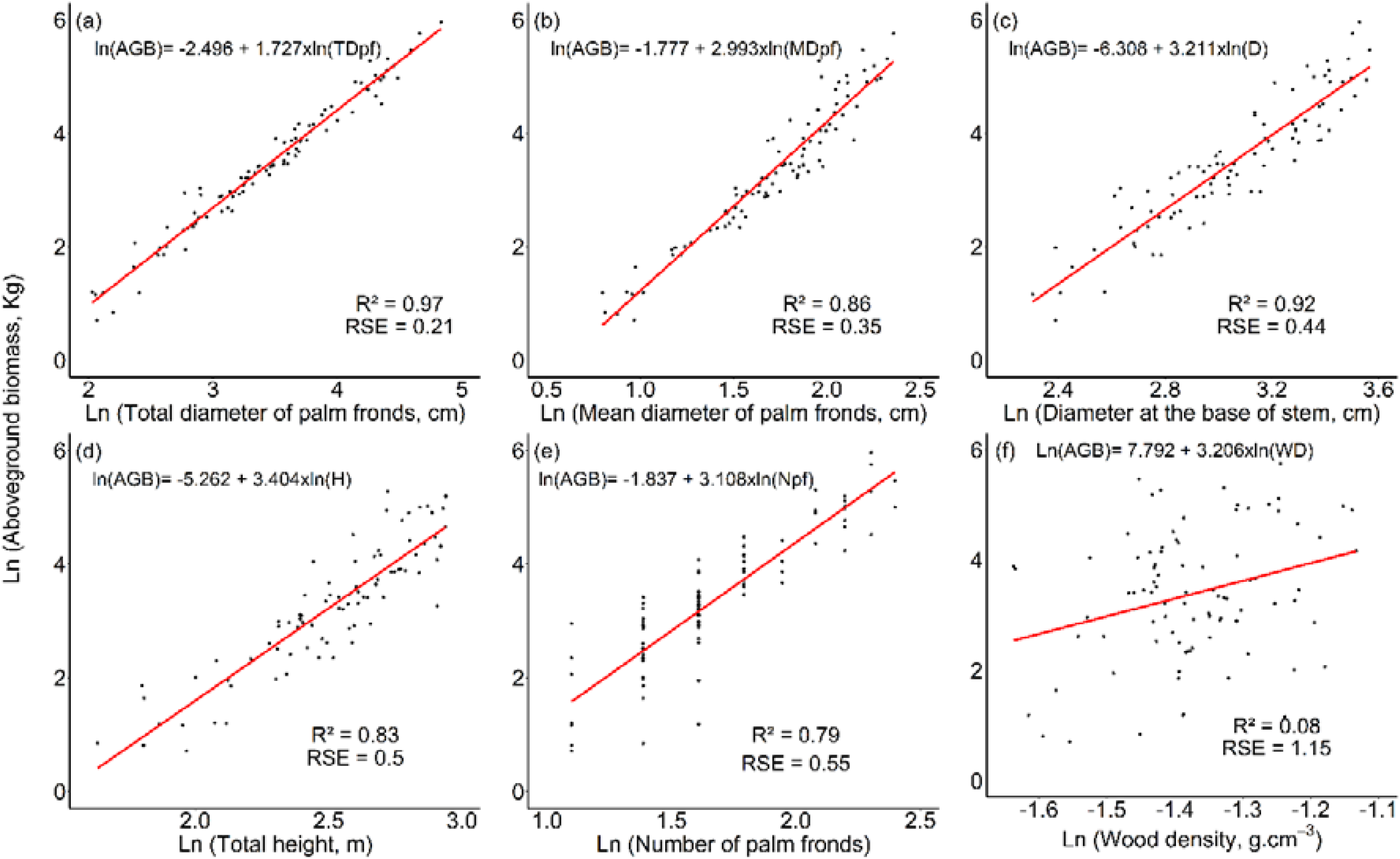
Allometric relationships between the aboveground biomass of *R. laurentii* stipe and each predictor variable.

Only *WD* showed a very weak relationship with *AGB*. The full list of equations is given in Table 2. Thus, when only one predictor variable was used, model 1 (m1), using *TD*_*pf*,_ was the best model (Table 2). Yet, the inclusion of other predictor variables (particularly *H*) significantly improved the goodness fit of the models, with the exception of *N*_*pf*_ or *WD* (Table 2). In fact, the combination of *TD*_*pf*_ with *H* increased the *R*^*2*^_*ad*j_ and significantly decreased the *AIC, RSE, RMSE* and *Bias*, compared to the models which used *TD*_*pf*_ in combination with *N*_*pf*_ (m5) or *WD* (m6). When three predictor variables were used in the linear model, the fit was slightly improved. The best *R*^*2*^_*adj*_, *AIC* and *RSE* values included the variables *TD*_*pf*_, *H* and *WD*, m11 in Table 2, in the following formula:

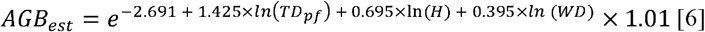

**Table 2.** Allometric equation fitted to estimate aboveground biomass (*AGB*: in kg of dry mass) of *Raphia laurentii* stipe in Congo Basin.

| Models |  | Coefficient and exponent estimate |  |  |  | Performance criteria |  |  |  |  | Diameter |
| --- | --- | --- | --- | --- | --- | --- | --- | --- | --- | --- | --- |
| N° | Formula | *a* | *b* | *c* | *d* | $R^2_{adj}$ | *RSE* | *AIC* | *RMSE* | *Bias* | range (n) |
| **m1** | $\ln(AGB) = a + b \times \ln(TD_{pf})$ | -2.496 | 1.727 | NA | NA | 0.97 | 0.21 | -19.48 | 15.44 | 0.02 | [2-15 cm |
| **m2** | $\ln(AGB) = a + b \times \ln(D)$ | -6,308 | 3,211 | NA | NA | 0,86 | 0,44 | 113,53 | 39.28 | -4.58 | |
| **m3** | $\ln(AGB) = a + b \times \ln(MD_{pf})$ | -1,777 | 2,993 | NA | NA | 0,92 | 0,35 | 69,6 | 28.77 | -5.59 | |
| **m4** | $\ln(AGB) = a + b \times \ln(H)$ | -5,262 | 3,404 | NA | NA | 0,82 | 0,5 | 136,13 | 38.51 | -5.55 | |
| **m5** | $\ln(AGB) = a + b \times \ln(N_{pf})$ | -1,837 | 3,108 | NA | NA | 0,79 | 0,55 | 151,96 | 37.73 | 8.61 | |
| **m6** | $\ln(AGB) = a + b \times \ln(WD)$ | 7,792 | 3,206 | NA | NA | 0,07 | 1,15 | 285,26 | 66.88 | 8.78 | |
| **m7** | $\ln(AGB) = a + b \times \ln(TD_{pf}) + c \times \ln(H)$ | -3.318 | 1.431 | 0.721 | NA | 0.98 | 0.18 | -46.25 | 15.37 | -1.54 | |
| **m8** | $\ln(AGB) = a + b \times \ln(TD_{pf}) + c \times \ln(N_{pf})$ | -2.431 | 2.087 | -0.769 | NA | 0.97 | 0.19 | -37.86 | 15.97 | -2.01 | |
| **m9** | $\ln(AGB) = a + b \times \ln(TD_{pf}) + c \times \ln(WD)$ | -1.751 | 1.707 | 0.493 | NA | 0.97 | 0.21 | -23.09 | 15.93 | 0.22 | |
| **m10** | $\ln(AGB) = a + b \times \ln(TD_{pf}) + c \times \ln(H) + d \times \ln(N_{pf})$ | -3.084 | 1.683 | 0.544 | -0.383 | 0.98 | 0.18 | -48.54 | 15.45 | -2.19 | |
| **m11** | $\ln(AGB) = a + b \times \ln(TD_{pf}) + c \times \ln(H) + d \times \ln(WD)$ | -2.691 | 1.425 | 0.695 | 0.395 | 0.98 | 0.18 | -49.15 | 15.71 | -1.55 | |
| **m12** | $\ln(AGB) = a + b \times \ln(TD_{pf})$ | -2.238 | 1.662 | NA | NA | 0.94 | 0.19 | -24.05 | 19.95 | -1.67 | [5-15 cm<br>[(60) |
| **m13** | $\ln(AGB) = a + b \times \ln(D)$ | -5.737 | 3.048 | NA | NA | 0.69 | 0.45 | 77.3 | 48.07 | -3.76 | |
| **m14** | $\ln(AGB) = a + b \times \ln(MD_{pf})$ | -2.573 | 3.4 | NA | NA | 0.77 | 0.39 | 59.84 | 33.89 | -5.59 | |
| **m15** | $\ln(AGB) = a + b \times \ln(H)$ | -5.027 | 3.349 | NA | NA | 0.64 | 0.48 | 86.29 | 45.59 | 0.27 | |
| **m16** | $\ln(AGB) = a + b \times \ln(N_{pf})$ | -0.272 | 2.353 | NA | NA | 0.8 | 0.36 | 50.73 | 41.86 | -2.59 | |
| **m17** | $\ln(AGB) = a + b \times \ln(WD)$ | 6.075 | 1.497 | NA | NA | 0.03 | 0.79 | 145.41 | 72.19 | -1.24 | |
| **m18** | $\ln(AGB) = a + b \times \ln(TD_{pf}) + c \times \ln(H)$ | -3.37 | 1.44 | 0.729 | NA | 0.96 | 0.17 | -38.32 | 18.41 | -1.25 | |
| **m19** | $\ln(AGB) = a + b \times \ln(TD_{pf}) + c \times \ln(N_{pf})$ | -2.478 | 1.964 | -0.491 | NA | 0.95 | 0.19 | -26.25 | 18.69 | -1.16 | |
| **m20** | $\ln(AGB) = a + b \times \ln(TD_{pf}) + c \times \ln(WD)$ | -1.92 | 1.653 | 0.209 | NA | 0.94 | 0.19 | -22.89 | 19.99 | -1.56 | |
| **m21** | $\ln(AGB) = a + b \times \ln(TD_{pf}) + c \times \ln(H) + d \times \ln(N_{pf})$ | -3.375 | 1.559 | 0.678 | -0.169 | 0.95 | 0.17 | -36.87 | 18.06 | -1.43 | |
| **m22** | $\ln(AGB) = a + b \times \ln(TD_{pf}) + c \times \ln(H) + d \times \ln(WD)$ | -3.232 | 1.44 | 0.718 | 0.08 | 0.95 | 0.17 | -36 | 18.44 | -1.24 | |

When only individuals with palm frond diameters greater than or equal to 5cm were used for model establishment, the *TD*_*pf*_ was the best predictor variable, followed by the *N*_*pf*_, *MD*_*pf*_, *D* and *H*. The *WD* was again the single worst predictor variable of the aboveground biomass. Thus, m16 can be used for an estimate of the biomass in the absence of the total diameter of the palm fronds. But when the *TD*_*pf*,_ *N*_*pf*_, *MD*_*pf*_, *D, H* and *WD* are available, the best model using more than one parameter is m18 (Table 2). However, m11 and m18 tend to underestimate *AGB* for largest *R. laurentii* individuals (Fig 6).

**Fig 6.**
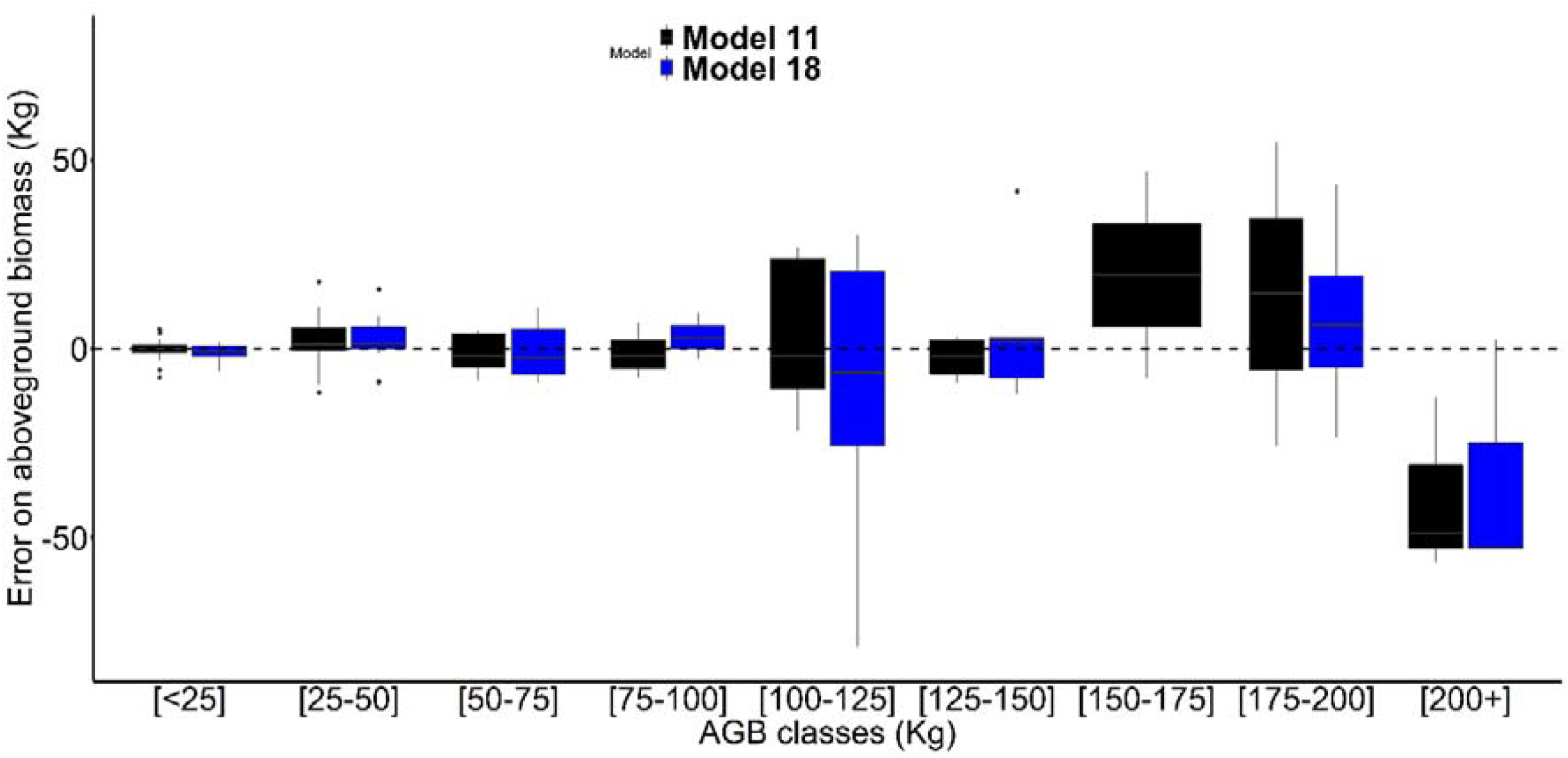
Goodness prediction of aboveground biomass (*AGB*) of model 11 and model 18. Mean relative error on *AGB* (***Error*** = [(***AGB***_***est***_ − ***AGB***_***obs***_)/***AGB***_***obs***_] × 100) for *AGB* classes.

The fastest and most cost-effective field method for estimating the *AGB* of *R. laurentii* stipes is counting palm fronds and using the single parameter equation, *N*_*pf*_, model 5 or model 16 in Table 2, for fronds ≥2 cm or ≥5 cm respectively. Therefore, the contribution of the AGB of *R. laurentii* stipes was assessed using model 16, on two 1 ha plots, the hardwood-dominated peat swamp forest EKG-02 and the palm-dominated peat swamp forest EKG-03. The contribution of *R. laurentii* individuals to the total above-ground biomass estimate was 8% and 41% for the tree-dominated EKG-02 and *R. laurentii* dominated EKG-03 plots, respectively. The percentage difference in biomass estimation between total above-ground biomass and above-ground biomass of trees was 9% and 71%, for the tree-dominated and *R. laurentii* dominated plot, respectively (Fig. 7).

**Fig 7.**
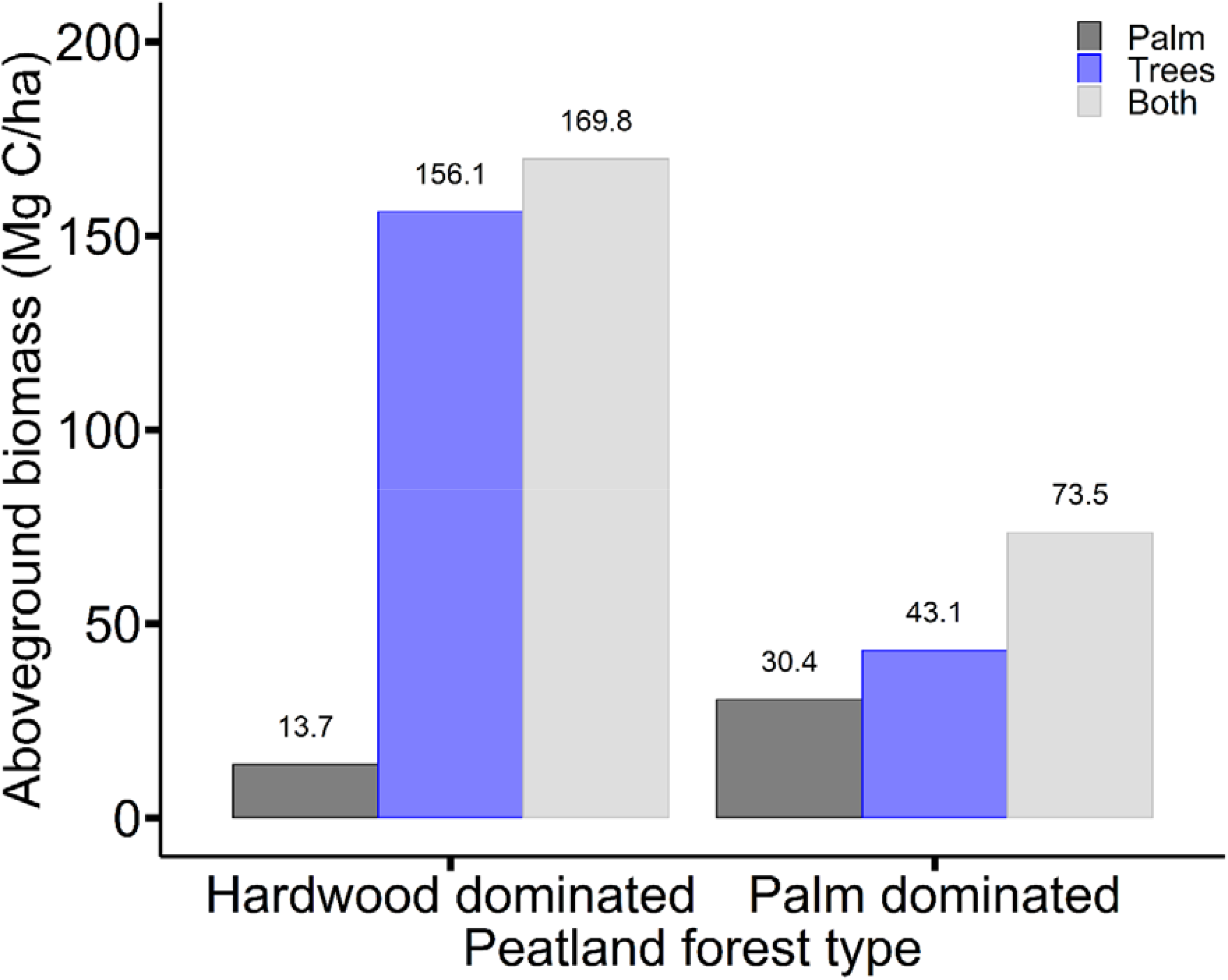
The contribution of the AGB of Raphia individuals in the total estimation of aboveground biomass carbon in the Congo Peatland.

## Discussion

### Aboveground biomass partition

Analysis of the distribution of aboveground biomass (*AGB*) between the different compartments of an individual of *R. laurentii* showed that the palm fronds represented at least 77 % of the total *AGB* in each diameter classes selected for the present study. This distribution of AGB between the stem and the palm fronds of *R. laurentii* was contrary to that observed for *Euterpe precatoria* Mart. (81 % in the stem) in the Amazonian Basin [10] and *Elaeis guineensis* Jacq. (73 % in the stem) in the Congo Basin [34]. This difference in biomass allocation can be linked to the architecture which differs from one type of palm to another [8,10]. Some palm species have a simple architecture, such as *Euterpe precatoria, Elaeis guineensis, Raphia hookeri* G. Mann and H. Wendl., *Cocos nucifera* L., *Hyphaene guineensis* Schumach. & Thonn., *Areca cotechu* L., which also makes it possible to estimate the stem height and the total palm height. Others, on the other hand, have a more complex architecture, such as *R. laurentii* and *Raphia monbuttorum* Drude [23]. For the latter, the presence of both living and dead fronds around the stem means that often the stem is not visible, impeding the measurement of its height. Practially, this means allometries which include frond measurements are superior to those including height. For *R. laurentii* the practical allometry must include the fronds. The distribution of biomass between sheath, petiole, rachis and leaflets also varies from one palm species to another. In this study, the biomass of the petioles and leaflets were greater than that of the rachis, which is contrary to the findings of [34] for *Elaeis guineensis*, where the biomass of the rachis (13.53%) was greater than that of the petioles (7.95%) and leaflets (6.02%). This difference can be attributed to the length and diameter of the petioles and also the consistency and number of pairs of leaflets which differ between palm species. For *Elaeis guineensis*, Sosef et al. [23] report petiole lengths of only 1– 1.25 m, with rachis lengths up to 8 m. Our results revealed that the length and diameter of the petioles of *R. laurentii* ranged from 0.07 – 8.26 m to 2.22–12.54 cm, respectively with a rachis length ranging from 0.43–12.58 m (Table 1). Thus the total length of a palm frond, including the petiole and the rachis is up to 19 m. The allows *R. laurentii* to contribute to the canopy in the swamp forests where it dominates, as there are relatively few trees (83 trees ≥10 cm diameter ha^-1^ in EKG-03), which are of modest size (mean diameter 10 cm, maximum 95.5 cm).

### Allometric equation for AGB estimation

We found that the best estimates of *R. laurentii AGB* were obtained when *TD*_*pf*_, *H* and *WD* were included as variables in the model (m11; Table 2). However, single variable models, using either *TD*_*pf*_, *MD*_*pf*_, *D, H or N*_*pf*_ also performed well when estimating *R. laurentii* AGB, with *TD*_*pf*_ as the best single variable to predict *AGB* (Table 2). *WD* was the only variable which was not suitable for predicting *R. laurentii AGB* in a single variable model. The finding that *TD*_*pf*_ was the best predictor variable of *R. laurentii AGB*, differs from the findings of other palm allometry studies and highlights the need for a *R. laurentii* specific allometric equation. For example both [18] and [11] found palm height was the best predictor variable of *AGB* for nine Amazonian palm species and for *Areca catechu* L. in Papua New Guinea, respectively. Whilst our results showed the inclusion of stem height improved *AGB* estimates, as a single predictor variable it was not the most suitable.

We therefore recommend, that when height data is available, m11, which uses *TD*_*pf*_, *H* and *WD* as variables, is used to estimate *R. laurnentii AGB*. Rather than sampling *WD* in the field, users of our allometric equation could use the *WD* values we provide in Table A1. However, owing to the tendency of m11 to underestimate the *AGB* of larger *R. laurentii* individuals, we recommend that m18, which includes the same variables but includes only stipes with palm frond diameters ≥ 5cm, is used for larger individuals.

The increase in AGB estimates for the two forest plots used in this study, particularly the palm-dominated plot, when estimates of *R. laurentii* biomass were included using m16, highlights the importance of including palms in aboveground biomass estimates. All the models we have developed here, (with exception of the simple linear *WD* model) permit us to estimate the AGB of *R. launrentii*, even when only very simple field measurements, such as the number of palm fronds per individual, are available.

If palm-rich peat swamp forest occupies 65,475 km^2^ of peatland in the central Congo Basin [4]. using our estimate of 30.4 Mg C ha^-1^ for the palm-dominated plot, we estimate that ∼1.99 million tonnes of carbon is stored aboveground in *R. launrentii* across the region. Prolific across the swamp forests of the Congo Basin, the inclusion of accurate estimations of *R. laurentii* palm biomass will be beneficial for Congo Basin countries seeking to benefit from results-based payments for REDD+ activities [35] and will allow a better estimate of forest carbon stocks and emissions across the region [3,36].

## Conclusion

We have developed the first allometric equation for the palm species *Raphia laurentii*, which permits the estimation of its *AGB*. Contrary to the findings of studies of other palm species, we find that the *AGB* of *R. laurentii* is concentrated mainly in the palm fronds (77 %), as it has a largely trunkless morphology. Subsequently, we find that the total diameter of palm fronds was the key variable to estimate *R. laurentii AGB*. This is different to the findings of allometry studies of other palm species, which have shown height to be the key predictor variable of palm *AGB*, highlighting the importance of a species specific allometric equation in the estimation of *R. launrentii AGB*. Whilst the total palm frond diameter single variable model gives good estimates of *AGB*, the model which includes height and wood density as variables gives the best estimates of *R. laurentii AGB*. The use of allometric models developed for the estimation of the biomass of *R. laurentii* stipes will allow a better estimation of the carbon stocks of the Congo Basin peat swamp forest. Future work should focus on other common palms in Africa, to improve biomass estimates and carbons stock estimates of these increasingly valuable ecosystems.

## Acknowledgements

We thank the villagers of Bolembe, Ekolongouma, Bethlem and Moungouma, for their assistance during the field data collection period. Thanks to the Ministries of the Environment, Forestry and Scientific Research of the Republic of Congo, Marien Ngouabi University and the Likouala governor for the research permits. We Thank the fieldworker team. We would also like to thank the University of Leeds (School of Geography) and the University of Edinburgh for providing the working environment during the scientific visit that enabled the finalisation of this manuscript. It is a pleasure to acknowledge Institut nationale de Recherche Forestière (IRF) for the provision of a third oven for drying the samples. At the end of this work, we would also like to thank Ifo S.A. and Helen Plante for logistical another support.

## Supporting information

**S1 Fig. Residue graphs of the model 11**

**S2 Fig. Residue graphs of the model 18**

**S1 Table.** Wood density values of the different compartments of an individual *Raphia laurentii*.

| Palm compartment | Mean palm diameter range (cm) | Wood density (g/cm<sup>3</sup>) |  |
| --- | --- | --- | --- |
|  |  | Mean | SD |
| Stem | [2 - 4] | 0,19 | 0,03 |
|  | [4 - 5] | 0,20 | 0,02 |
|  | [5 - 6] | 0,21 | 0,04 |
|  | [6 - 7] | 0,19 | 0,03 |
|  | [7 - 8] | 0,22 | 0,04 |
|  | [+8] | 0,27 | 0,05 |
| Sheath | [2 - 4] | 0,27 | 0,05 |
|  | [4 - 5] | 0,26 | 0,03 |
|  | [5 - 6] | 0,26 | 0,04 |
|  | [6 - 7] | 0,27 | 0,04 |
|  | [7 - 8] | 0,25 | 0,04 |
|  | [+8] | 0,25 | 0,04 |
| Petiole | [2 - 4] | 0,22 | 0,03 |
|  | [4 - 5] | 0,21 | 0,02 |
|  | [5 - 6] | 0,21 | 0,03 |
|  | [6 - 7] | 0,21 | 0,02 |
|  | [7 - 8] | 0,20 | 0,03 |
|  | [+8] | 0,20 | 0,02 |
| Rachis | [2 - 4] | 0,27 | 0,04 |
|  | [4 - 5] | 0,27 | 0,02 |
|  | [5 - 6] | 0,29 | 0,02 |
|  | [6 - 7] | 0,29 | 0,04 |
|  | [7 - 8] | 0,28 | 0,04 |
|  | [+8] | 0,29 | 0,04 |

## Notes

### Competing Interest Statement

The authors have declared no competing interest.

## References

1. Brown S, Iverson LR. Biomass estimates for tropical forests. World Resource Review. 1992;4: 366–384.

2. Unfcc C. Rapport de la Conférence des Parties sur sa seizième session, tenue à Cancún du 29 novembre au 10 décembre 2010. 2010 pp. 1–34. Report No.: FCCC/CP/2010/7/Add.1.

3. Angelsen A, Brockhaus M, Sunderlin WD, Verchot LV. Analyse de la REDD+C: Les enjeux et les choix. Arild Angelsen. Bogor, Indonésie: CIFOR; 2013.

4. Dargie GC, Lewis SL, Lawson IT, Mitchard ETA, Page SE, Bocko YE, et al. Age, extent and carbon storage of the central Congo Basin peatland complex. Nature. 2017;542: 86–90. doi:10.1038/nature21048

5. Bocko YE, Ifo SA, Loumeto JJ. Quantification Des Stocks De Carbone De Trois Pools Clés De Carbone En Afrique CentraleC: Cas De La Forêt Marécageuse De La Likouala (Nord Congo). Eur Sci J ESJ. 2017;13: 438. doi:10.19044/esj.2017.v13n5p438

6. Hubau W, Lewis SL, Phillips OL, Affum-Baffoe K, Beeckman H, Cuní-Sanchez A, et al. Asynchronous carbon sink saturation in African and Amazonian tropical forests. Nature. 2020;579: 80–87. doi:10.1038/s41586-020-2035-0

7. Muscarella R, Emilio T, Phillips OL, Lewis SL, Slik F, Baker WJ, et al. The global abundance of tree palms. McGeoch M, editor. Glob Ecol Biogeogr. 2020;29: 1495–1514. doi:10.1111/geb.13123

8. Cole TG, Ewel JJ. Allometric equations for four valuable tropical tree species. For Ecol Manag. 2006;229: 351–360. doi:10.1016/j.foreco.2006.04.017

9. Goodman RC, Phillips OL, del Castillo Torres D, Freitas L, Cortese ST, Monteagudo A, et al. Amazon palm biomass and allometry. For Ecol Manag. 2013;310: 994–1004. doi:10.1016/j.foreco.2013.09.045

10. Da Silva F, Suwa R, Kajimoto T, Ishizuka M, Higuchi N, Kunert N. Allometric Equations for Estimating Biomass of Euterpe precatoria, the Most Abundant Palm Species in the Amazon. Forests. 2015;6: 450–463. doi:10.3390/f6020450

11. Prayogo C, Sari RR, Asmara DH, Rahayu S, Hairiah K. Allometric Equation for Pinang (Areca catechu) Biomass and C Stocks. AGRIVITA J Agric Sci. 2018;40. doi:10.17503/agrivita.v40i3.1124

12. Lewis K, Rumpang E, Kho LK, McCalmont J, Teh YA, Gallego-Sala A, et al. An assessment of oil palm plantation aboveground biomass stocks on tropical peat using destructive and non-destructive methods. Sci Rep. 2020;10: 2230. doi:10.1038/s41598-020-58982-9

13. Dargie GC, Lawson IT, Rayden TJ, Miles L, Mitchard ETA, Page SE, et al. Congo Basin peatlands: threats and conservation priorities. Mitig Adapt Strateg Glob Change. 2018;24: 669–686. doi:10.1007/s11027-017-9774-8

14. Picard et al. Organisation des Nations Unies pour l’alimentation et l’agriculture. Popul Fr Ed. 2012;5: 764. doi:10.2307/1523706

15. Chave J, Andalo C, Brown S, Cairns MA, Chambers JQ, Eamus D, et al. Tree allometry and improved estimation of carbon stocks and balance in tropical forests. Oecologia. 2005;145: 87–99. doi:10.1007/s00442-005-0100-x

16. Loubota Panzou GJ, Fayolle A, Jucker T, Phillips OL, Bohlman S, Banin LF, et al. Pantropical variability in tree crown allometry. Kerkhoff A, editor. Glob Ecol Biogeogr. 2021;30: 459–475. doi:10.1111/geb.13231

17. Avalos G, Fernández Otárola M. Allometry and stilt root structure of the neotropical palm Euterpe precatoria (Arecaceae) across sites and successional stages. Am J Bot. 2010;97: 388–394. doi:10.3732/ajb.0900149

18. Goodman RC, Phillips OL, del Castillo Torres D, Freitas L, Cortese ST, Monteagudo A, et al. Amazon palm biomass and allometry. For Ecol Manag. 2013;310: 994–1004. doi:10.1016/j.foreco.2013.09.045

19. Davenport IJ, McNicol I, Mitchard ETA, Dargie G, Suspense I, Milongo B, et al. First Evidence of Peat Domes in the Congo Basin using LiDAR from a Fixed-Wing Drone. Remote Sens. 2020;12: 2196. doi:10.3390/rs12142196

20. ANAC. Données Météorologique de la région de la Likouala – Données météorologiques du district d’Impfondo pour les 50 dernières années. Base de données annuelles de l’agence nationale de l’aviation civile (ANAC), Ministère de transport et de l’aviation civile, Brazzaville, Congo. 2020.

21. Rainey HJ, Iyenguet FC, Malanda G-AF, Madzoké B, Santos DD, Stokes EJ, et al. Survey of Raphia swamp forest, Republic of Congo, indicates high densities of Critically Endangered western lowland gorillas Gorilla gorilla gorilla. Oryx. 2010;44: 124. doi:10.1017/S003060530999010X

22. Bocko YE, Dargie G, Yoka J. Répartition spatiale de la richesse floristique des forêts marécageuses de la Likouala, Nord-Congo. 2016; 13.

23. Sosef MSM, Florence J, Ngok Banak L, Bourobou Bourobou HP, Bissiengou P. Flore du Gabon RaphiaC: Palmae. 2019.

24. Avalos G, Sylvester O. Allometric estimation of total leaf area in the neotropical palm Euterpe oleracea at La Selva, Costa Rica. Trees. 2010;24: 969–974. doi:10.1007/s00468-010-0469-y

25. Walker SM, Pearson TR, Casarim FM, Harris N, Petrova S, Grais A, et al. Standard Operating Procedures for Terrestrial Carbon Measurement. 2012; 96.

26. Fayolle A, Ngomanda A, Mbasi M, Barbier N, Bocko Y, Boyemba F, et al. A regional allometry for the Congo basin forests based on the largest ever destructive sampling. For Ecol Manag. 2018;430: 228–240. doi:10.1016/j.foreco.2018.07.030

27. Vahedi AA, Mataji A, Babayi-Kafaki S, Eshaghi-Rad J, Hodjati SM, Djomo A. Allometric equations for predicting aboveground biomass of beech-hornbeam standsin the Hyrcanian forests of Iran. J For Sci. 2014;60: 236–247. doi:10.17221/39/2014-JFS

28. Fonton NH, Medjibé V, Djomo A, Kondaoulé J, Rossi V, Ngomanda A, et al. Analyzing Accuracy of the Power Functions for Modeling Aboveground Biomass Prediction in Congo Basin Tropical Forests. Open J For. 2017;07: 388–402. doi:10.4236/ojf.2017.74023

29. Chave J, Réjou-Méchain M, Búrquez A, Chidumayo E, Colgan MS, Delitti WBC, et al. Improved allometric models to estimate the aboveground biomass of tropical trees. Glob Change Biol. 2014;20: 3177–3190. doi:10.1111/gcb.12629

30. Parresol B. Assessing Tree and Stand Biomass. A Review with Examples and Critical Comparisons. For Sci. 1999; 573–593.

31. Manuri S, Brack C, Noor’an F, Rusolono T, Anggraini SM, Dotzauer H, et al. Improved allometric equations for tree aboveground biomass estimation in tropical dipterocarp forests of Kalimantan, Indonesia. For Ecosyst. 2016;3: 28. doi:10.1186/s40663-016-0087-2

32. Conti G, Gorné LD, Zeballos SR, Lipoma ML, Gatica G, Kowaljow E, et al. Developing allometric models to predict the individual aboveground biomass of shrubs worldwide. Kerkhoff A, editor. Glob Ecol Biogeogr. 2019;28: 961–975. doi:10.1111/geb.12907

33. Balima LH, Nacoulma BMI, Bayen P, Dimobe K, Kouamé FN, Thiombiano A. Aboveground biomass allometric equations and distribution of carbon stocks of the African oak (Afzelia africana Sm.) in Burkina Faso. J For Res. 2020;31: 1699–1711. doi:10.1007/s11676-019-00955-4

34. Migolet P, Goïta K, Ngomanda A, Mekui Biyogo AP. Estimation of Aboveground Oil Palm Biomass in a Mature Plantation in the Congo Basin. Forests. 2020;11: 544. doi:10.3390/f11050544

35. FAO. Considérations techniques relatives à l’établissement de niveaux d’émissions de référence pour les forêts et/ou niveaux de référence pour les forêts dans le contexte de la REDD+ au titre de la CCNUCC. 2015.

36. Rozendaal DMA, Requena Suarez D, De Sy V, Avitabile V, Carter S, Adou Yao CY, et al. Aboveground forest biomass varies across continents, ecological zones and successional stages: refined IPCC default values for tropical and subtropical forests. Environ Res Lett. 2022;17: 014047. doi:10.1088/1748-9326/ac45b3

