## Supplementary figures and images for "Allometric equation for the commonest palm in the Central Congo Peatlands, *Raphia laurentii* De Wild"

### S1_Fig.tif.docx

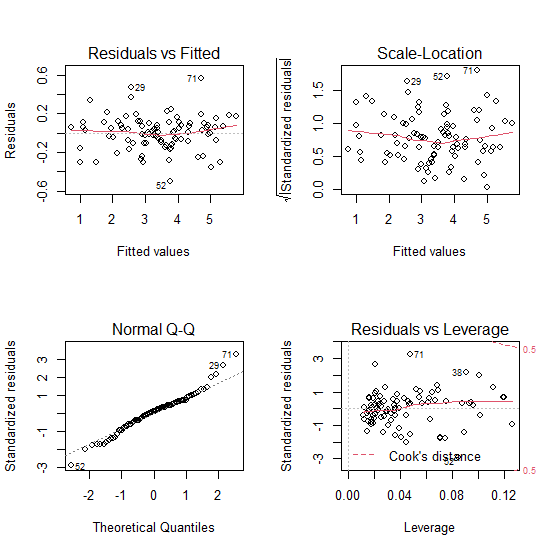

S1 Fig. Residue graphs of the model 11

### S2_Fig.tif.docx

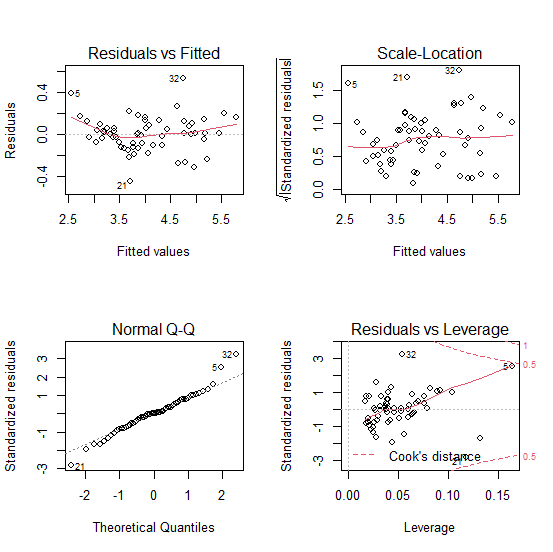

S2 Fig. Residue graphs of the model 18
